# Selection-driven cost-efficiency optimisation of transcripts modulates gene evolutionary rate in bacteria

**DOI:** 10.1101/136861

**Authors:** Emily A. Seward, Steven Kelly

## Abstract

**Background:** Most amino acids are encoded by multiple synonymous codons. However synonymous codons are not used equally and this biased codon use varies between different organisms. It has previously been shown that both selection acting to increase codon translational efficiency and selection acting to decrease codon biosynthetic cost contribute to differences in codon bias. However, it is unknown how these two factors interact or how they affect molecular sequence evolution.

**Results:** Through analysis of 1,320 bacterial genomes we show that bacterial genes are subject to multi-objective selection-driven optimisation of codon use. Here, selection acts to simultaneously decrease transcript biosynthetic cost and increase transcript translational efficiency, with highly expressed genes under the greatest selection. This optimisation is not simply a consequence of the more translationally efficient codons being less expensive to synthesise. Instead, we show that tRNA gene copy number alters the cost-efficiency trade-off of synonymous codons such that for many species such that selection acting on transcript biosynthetic cost and translational efficiency act in opposition. Finally, we show that genes highly optimised to reduce cost and increase efficiency show reduced rates of synonymous and non-synonymous mutation.

**Conclusions:** This analysis provides a simple mechanistic explanation for variation in evolutionary rate between genes that depends on selection-driven cost-efficiency optimisation of the transcript. These findings reveal how optimisation of resource allocation to mRNA synthesis is a critical factor that determines both the evolution and composition of genes.

## Background

Production of proteins is a primary consumer of cell resources [1]. It requires allocation of cellular resources to production of RNA sequences as well as allocation of resources to production of nascent polypeptide chains. Whilst a protein’s amino acid sequence is functionally constrained, redundancy in the genetic code means that multiple nucleotide sequences can code for the same protein. Since the biosynthetic cost and translational efficiency of synonymous codons varies, biased use of synonymous codons makes it possible to reduce the expenditure of cellular resources on mRNA production without altering the encoded protein sequence. Thus, it is possible to reduce resource allocation to protein synthesis without altering the encoded protein or affecting protein abundance. This is done by reducing transcript sequence cost or by increasing the efficiency with which those transcripts can be translated into protein. Consistent with this, it has been demonstrated that natural selection acts both to reduce biosynthetic cost of RNA sequences [2,3], and to increase the efficiency with which those RNA sequences can template the encoded polypeptide chain [4–10]. However, though selection has been shown to act on codon biosynthetic cost and translational efficiency independently, it is unknown how these two factors interact or whether optimisation of one factor inherently results in optimisation of the other. It should be noted that in addition to factors acting on resource allocation, functional constraints are also known to bias patterns of codon use, for example, RNA structural constraints to facilitate thermal adaptation and translational initiation [11–13], RNA sequence constraints to preserve splice sites [14], and translational constraints to ensure accurate protein folding [15–17]. However, since those factors primarily act on individual sites or sets of sites within genes and are independent of resource allocation, they were not considered further in this analysis.

Different species employ different strategies to decode synonymous codons [18]. These strategies make use of ‘wobble’ base pairing between the 3^rd^ base of the codon and the 1^st^ base of the anticodon to facilitate translation of all 61 sense codons using a reduced set of tRNAs. As the translational efficiency of a codon is a function of the number of tRNAs that can translate that codon, and as different species encode different subsets of tRNA genes, the same codon is not necessarily equally translationally efficient in all species. In contrast, the biosynthetic cost of a codon of RNA is determined by the number and type of atoms contained within that codon and the number of high energy phosphate bonds required for their assembly. As translational efficiency varies between species but biosynthetic cost does not, it was hypothesised that this must create a corresponding variation in the codon cost-efficiency trade-off between species. For example biosynthetically cheap codons might be translationally efficient in one species but inefficient in another. We further hypothesised that variation in the codon cost-efficiency trade-off would limit the extent to which a transcript could be optimised to be both biosynthetically inexpensive and translationally efficient.

Here, we show that natural selection acts genome-wide to reduce cellular resource allocation to mRNA synthesis by solving the multi-objective optimisation problem of minimising transcript biosynthetic cost whilst simultaneously maximising transcript translational efficiency. We show that this optimisation is achieved irrespective of the codon cost-efficiency trade-off of a species, and that the extent to which resource allocation is optimised is a function of the production demand of that gene. Finally, we reveal that selection-driven optimisation of resource allocation provides a novel mechanistic explanation for differences in evolutionary rates between genes, and for the previously unexplained correlation in synonymous and non-synonymous mutation rates of genes.

## Results

### Selection acts to reduce biosynthetic cost and increase translational efficiency of transcript sequences

Although selection has been shown to reduce resource allocation to mRNA production by reducing codon biosynthetic cost or increasing translational efficiency independently [2–10], it is unknown how these two factors interact or whether optimisation of one factor inherently results in optimisation of the other. To address this, an analysis was conducted on 1,320 bacterial species representing 730 different genera to establish if they were either under selection to increase codon translational efficiency, reduce codon biosynthetic cost or a combination of the two (Table S1). For each species, genome-wide values for mutation bias towards GC [M_b_], selection on transcript translational efficiency [*S*_*t*_] and selection on transcript biosynthetic cost [*S_c_*] were inferred (Fig. 1). This was done using the complete set of open reading frames and tRNAs encoded in that species’ genome using the SK model [2] implemented using CodonMuSe (see Methods). Genome-wide GC content varied from 26% to 75% and so encompassed almost the entire range of known bacterial genome GC values ^[19]^, this large variation in content was reflected in the range of values observed for M_b_ (Fig. 1a, mean = 0.44). Of the 1,320 species in this analysis, 91% had negative *S*_*c*_ values (mean *S*_*c*_ = −0.08), indicating a genome-wide selective pressure to reduce the biosynthetic cost of transcript sequences through biased synonymous codon use (Fig. 1b). This observation is consistent with previous studies that revealed analogous effects when nitrogen or energy were limited [2,3]. Similarly, 78% of species had positive values for *S*_*t*_ (mean *S*_*t*_ = 0.1), indicating a genome-wide selective pressure to increase the translational efficiency of transcript sequences (Fig. 1c). This is consistent with multiple examples where a strong pressure has been shown to favour high translational efficiency [4–10]. Moreover, 74% of species had both a negative *S*_*c*_ value and a positive *S*_*t*_ value, demonstrating that selection is not mutually exclusive when acting on translational efficiency and codon biosynthetic cost. Indeed, the majority of species experience selection to reduce transcript biosynthetic cost while simultaneously maximising transcript translational efficiency.

**Fig 1.**
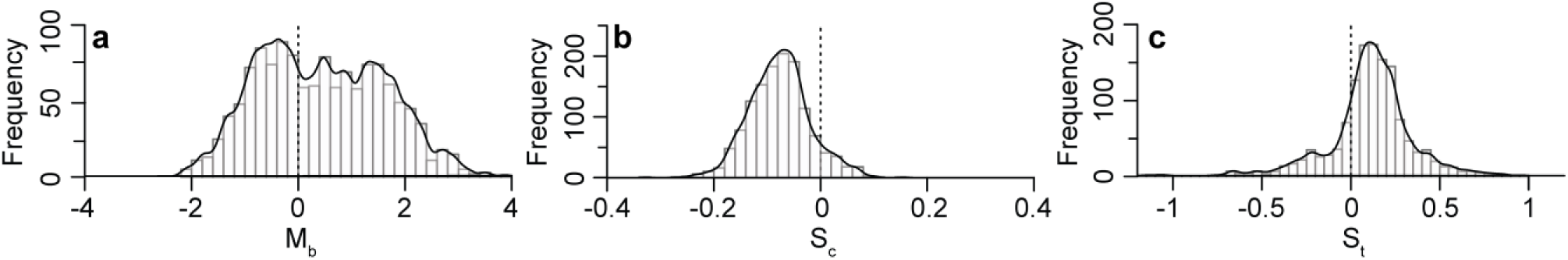
Bacterial genomes show selection to reduce nucleotide cost (-S_c_) and increase translational efficiency (+S_t_) Genome-wide values for 1,320 bacterial species covering 730 genera for **a)** mutation bias towards GC (M_b_). Positive values indicate mutation bias towards GC. Negative values indicate mutation bias towards AT. **b)** Strength of selection acting on codon biosynthetic cost (S_c_). Negative values indicate selection acting to reduce biosynthetic cost. **c)** Strength of selection acting on codon translational efficiency (S_t_). Positive values indicate selection acting to increase codon translational efficiency.

### More translationally efficient bacterial codons are generally more biosynthetically costly

The biosynthetic cost of a codon can be defined as the number and type of atoms contained within the codon or the number of high energy phosphate bonds required for their assembly. Natural selection acting on biosynthetic cost, both in terms of nitrogen atoms [2] or energetic requirements [3], has been shown to play a role in promoting biased patterns of synonymous codon use. However, as the energy and nitrogen cost of a codon correlate almost perfectly (Fig. 2a), it is not possible to distinguish which factor is responsible for biased patterns of codon use in the absence of additional information about the biology of the organism in question. Nonetheless, given the near perfect correlation, analysis of selection acting on overall codon biosynthetic cost can be approximated by analysis of either nitrogen or energetic requirements.

**Fig. 2.**
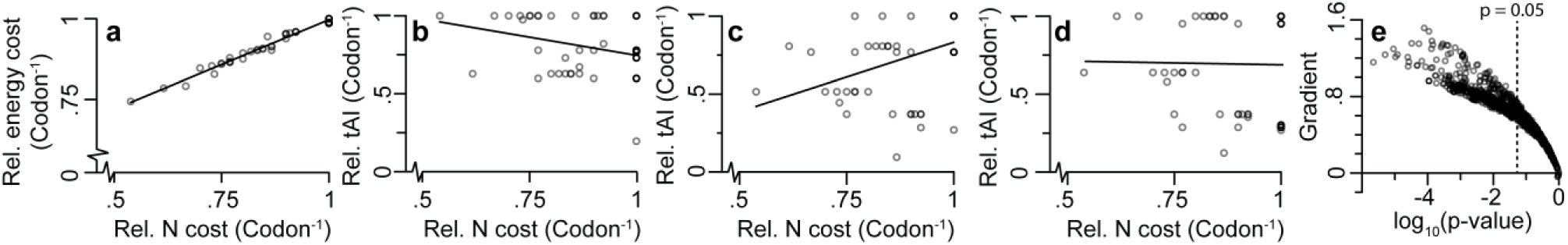
Different tRNA sparing strategies alter a species’ codon cost-efficiency trade-off. Codon nitrogen cost (N cost) correlates almost perfectly with codon energetic cost (p < 0.05, y = 0.6x + 0.44, R^2^ = 0.98). **b)** A full complement of tRNAs has a negative correlation between codon biosynthetic cost and translational efficiency (tAI) (p< 0.05, y = −0.5x + 1.21, R^2^ = 0.10). **c)** tRNA sparing strategy 1 (NNU codons translated by GNN anticodons) has a positive correlation between codon biosynthetic cost and translational efficiency (p<0.05, y = 0.9x – 0.06, R^2^ = 0.18). **d)** tRNA sparing strategy 2 (strategy 1 + NNG codons translated by UNN anticodons) has no significant correlation between codon biosynthetic cost and translational efficiency (p > 0.05, y = 0.74, R^2^ = 0). **e)** None of the 1,320 bacterial species in this analysis have a significant negative correlation between codon cost and translational efficiency (p > 0.05). The y-axis is the gradient of the line of best fit between codon biosynthetic cost and translational efficiency.

Codon translational efficiency is generally measured using the tRNA adaptation index (tAI), which considers both the abundance of iso-accepting tRNAs and wobble-base pairing [20]. Since tRNA gene copy number varies between species, there is a corresponding variation in the relative translational efficiency of their associated codons [18,21]. Therefore, the relationship between codon biosynthetic cost and codon translational efficiency (referred to from here on as the codon cost-efficiency trade-off) must vary between species. For example, a hypothetical species encoding a full complement of tRNAs, each present as a single copy, would have a negative correlation between cost and efficiency (Fig. 2b). In contrast, a hypothetical species that employed tRNA sparing strategy 1 (no ANN tRNAs) or strategy 2 (no ANN or CNN tRNAs) [18], would show a positive (Fig. 2c) or no (Fig. 2d) correlation between cost and efficiency respectively. Therefore, a broad range of codon cost-efficiency trade-offs is possible and the gradient of this trade-off is dependent on the tRNA gene copy number of a given species.

None of the 1,320 species used in this analysis contained a full complement of tRNAs. Moreover, only two species strictly adhered to a single sparing strategy for all synonymous codon groups (e.g. *Escherichia coli* uses strategy 2 for decoding alanine but strategy 1 for decoding glycine). Given that neither tRNA sparing strategy 1 nor 2 led to a negative correlation between cost and efficiency, it is therefore expected that species would have either a positive or no correlation between codon cost and efficiency. Furthermore, given the many different potential tRNA complements, it is anticipated that a continuum of gradients in trade-off between cost and efficiency would be observed. To assess this, the codon cost-efficiency trade-off was calculated for the 1,320 bacterial species (Fig. 2e). As expected, species with a significant negative correlation between cost and efficiency were not observed. Instead, all species exhibited either positive or non-significant correlations between codon cost and efficiency (Fig. 2e). Thus in general, the synonymous codons that are most translationally efficient are those that consume the most resources for biosynthesis.

### Genes that experience the strongest selection for increased transcript translational efficiency are also under the strongest selection to reduce biosynthetic cost

Given that the majority of species exhibited selection to reduce cost and increase translational efficiency at the genome-wide level, the extent to which this was also seen at the level of an individual gene within species was determined. Here, the strength of selection acting on transcript translational efficiency and strength of selection on transcript biosynthetic cost were inferred for each individual gene in each species. The relationship between *S*_*c*_ and *S*_*t*_ was then compared for each species. For example in *Escherichia coli,* which doesn’t have a strong cost-efficiency trade-off, there is a significant negative correlation between *S*_*c*_ and *S*_*t*_ (Fig. 3a). Here, the genes that experienced the greatest selection to increase efficiency are those that experienced the greatest selection to reduce biosynthetic cost. The same phenomenon was also observed for *Lactobacillus amylophilus,* a species with a strong codon cost-efficiency trade-off (Fig. 3b). Overall, significant correlations between *S*_*c*_ and *S*_*t*_ for individual genes were observed for 91% of species (p < 0.05, Fig. 3c). Therefore irrespective of the codon cost-efficiency trade-off, selection is performing multi-objective optimisation of transcript sequences to reduce their biosynthetic cost while increasing their translational efficiency and thereby reducing resource allocation to mRNA production.

**Fig. 3.**
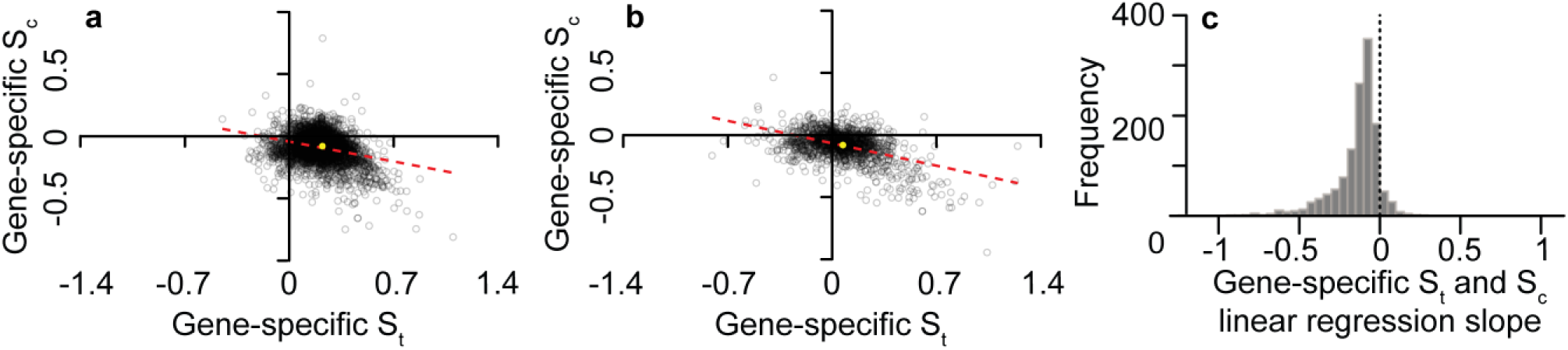
The genes under the strongest selection for translational efficiency (+S_t_) are also under the strongest selection to reduce nucleotide cost (-S_c_) Scatterplots of gene-specific St and Sc values for **a)** *Escherichia coli* **b)** *Lactobacillus amylophilus*. In both cases the line of best fit is shown (red) and the yellow dot is the genome-wide best-fit value for each species. Each point has been set to an opacity of 20% so density can be judged. **c)** Histogram of the slope between S_c_ and S_t_ for individual genes for each of the 1,320 bacterial species in this analysis.

As the most highly expressed genes in a cell comprise the largest proportion of cellular RNA, the strength of selection experienced by a gene is thought to be dependent on the mRNA abundance of that gene [22–24]. In agreement with this, evaluation of the relative mRNA abundance of genes in *E. coli* revealed that the most highly expressed genes exhibited the greatest selection to reduce transcript biosynthetic cost (Fig. 4a) whilst also showing the strongest selection to increase transcript translational efficiency (Fig. 4b). Thus, selection acts in proportion to relative mRNA abundance to perform multi-objective optimisation of codon bias in order to reduce resource allocation to transcript sequences through production of low cost, high efficiency transcripts.

**Fig. 4.**
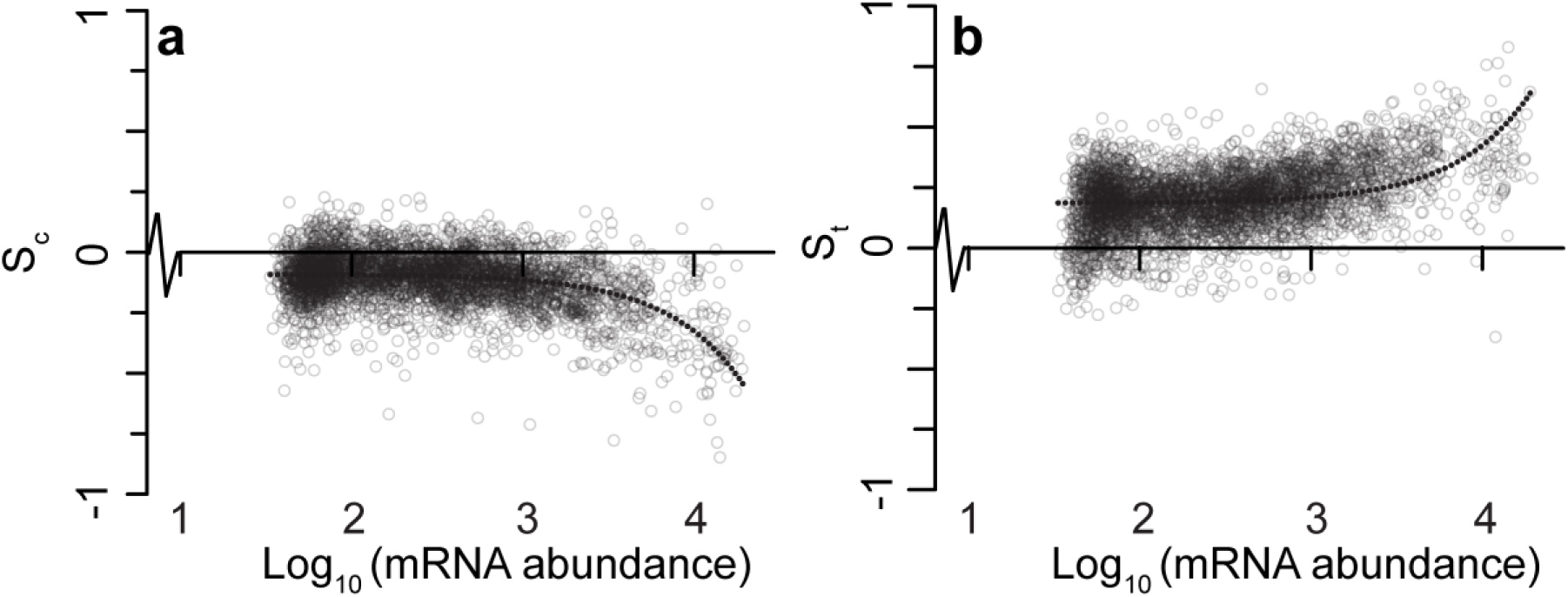
Selection acts in proportion to mRNA abundance to decrease codon biosynthetic cost and increase codon translational efficiency in *Escherichia coli*. **a)** There is a negative correlation between selection acting on codon biosynthetic cost (*S*_*c*_) and mRNA abundance. The linear line of best fit (shown here on a log scale) has an R^2^ value of 0.18. **b)** There is a positive correlation between selection acting to increase codon translational efficiency (*S*_*t*_) and gene expression. The linear line of best fit (shown here on a log scale) has an R^2^ value of 0.13. Each point has been set to an opacity of 20% so density can be judged.

### Sequence optimisation for cost and efficiency constrains molecular evolution rate

Given that codon choice has been shown to provide a selective advantage per codon per generation [25], it was hypothesised that the extent to which a transcript is jointly optimised for codon cost and efficiency would constrain the rate at which the underlying gene sequence can evolve. Specifically, the more highly optimised a transcript is for both biosynthetic cost and translational efficiency, the higher the proportion of spontaneous mutations that would reduce the cost-efficiency optimality of the transcript sequence. Therefore, spontaneous mutations in highly optimised genes are more likely to be deleterious than spontaneous mutations in less optimised genes. As deleterious mutations are lost more rapidly from the population than neutral mutations, the more highly optimised a gene sequence is, the lower its apparent evolutionary rate should be.

To test this hypothesis the complete set of gene sequences from *E.coli* was subject to stochastic *in silico* mutagenesis and the proportion of single nucleotide mutations that resulted in reduced transcript cost-efficiency optimality was evaluated. As expected, the proportion of deleterious mutations increased linearly with transcript sequence optimality. This effect was seen for both synonymous (Fig. 5a) and non-synonymous mutations (Fig. 5b). The effect in non-synonymous mutations is seen because a single base mutation from an optimal codon encoding one amino acid is unlikely to arrive at an equally optimal (or better) codon encoding any other amino acid. Thus as expected, the more optimal a codon is, the less likely a spontaneous mutation will result in a codon with higher optimality irrespective of whether that codon encodes the same amino acid.

**Fig. 5.**
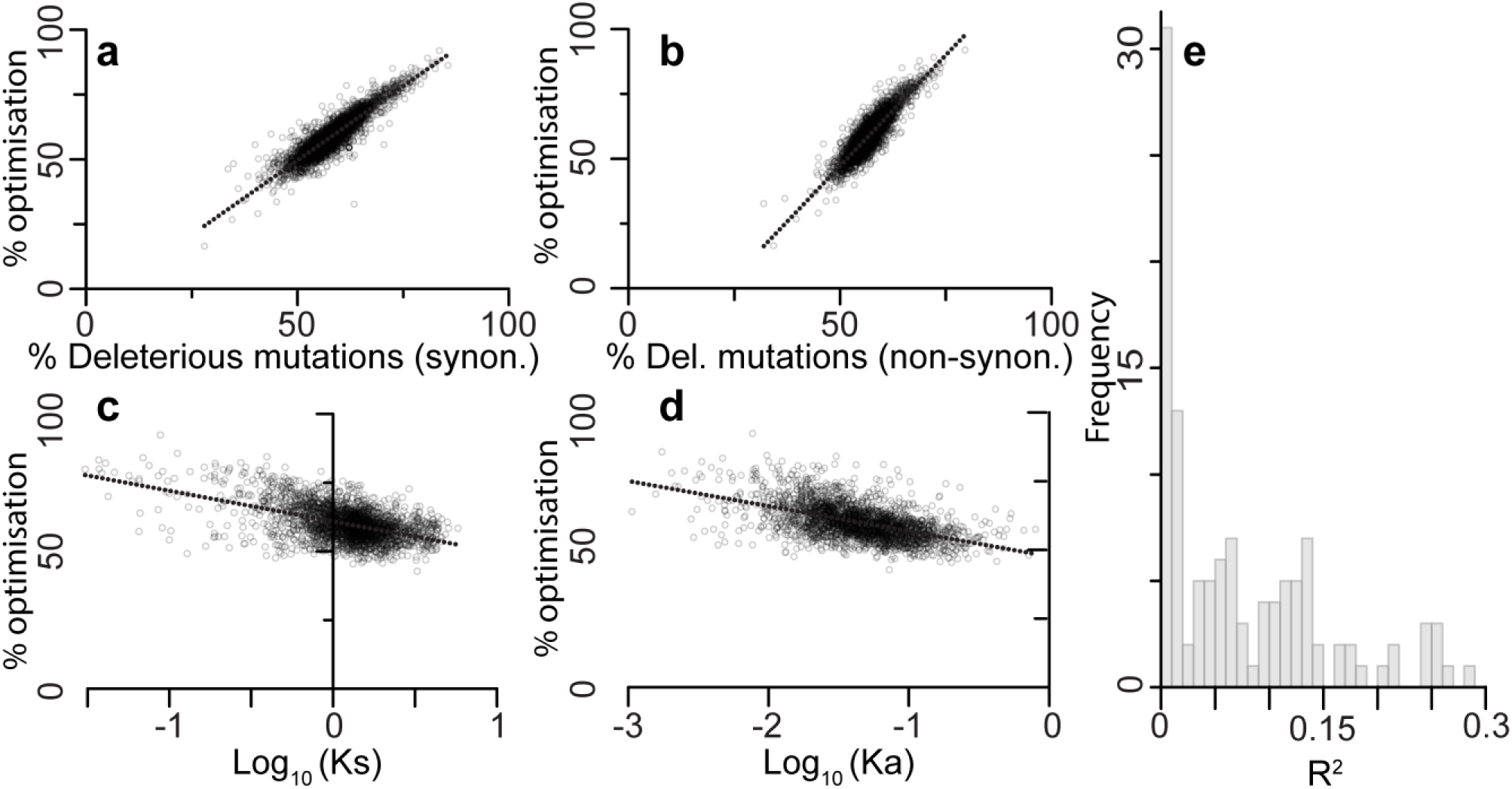
Selection-driven optimisation of resource allocation is a critical factor that determines molecular evolution rate. Highly cost-efficiency optimised genes have a higher proportion of deleterious **a)** synonymous (y = 1.15x −8, R^2^ = 0.81) and **b)** non-synonymous (y = 1.71x −38, R^2^ = 0.78) mutations. Orthologous genes in *Escherichia coli* and *Salmonella enterica* show a negative correlation between sequence cost-efficiency optimisation and the rate of **c)** synonymous mutations (*K*_*s*_) (y = −11x + 61, R^2^ = 0.26) and **d)** non-synonymous mutation (*K*_*a*_) (y =-9x + 48, R^2^ = 0.28). **e)** histogram of proportion of gene evolutionary rate explained by selection-driven cost-efficiency optimisation of transcript sequences.

The extent to which transcript sequences in *E. coli* were jointly cost-efficiency optimised was compared to the synonymous (*K*_*s*_) and non-synonymous (*K*_*a*_) mutation rate of that gene, estimated from comparison with *Salmonella enterica*. Consistent with the hypothesis, the rate of synonymous (*k*_*s*_ Fig. 5c) and non-synonymous (*k*_*a*_ Fig. 5d) changes were directly proportional to the extent to which the gene sequence had been optimised by natural selection for low biosynthetic cost and high translational efficiency (Fig. 5a and b). While efficiency optimisation explained more of the variance in gene evolutionary rate, the linear regression model that considered both cost and efficiency optimisation was significantly better than models that considered either factor alone, whether or not derived optimisation values or raw tAI and biosynthetic costs were considered (Supplementary Fig. S1, ANOVA, p < 0.001). Therefore, this analysis provides a mechanistic explanation for previous studies that found a strong correlation between non-synonymous evolutionary rate and mRNA abundance [22]. To determine if this relationship was also observed for other bacteria, an additional 177 species-pairs were analysed (Fig. 5e). Of these species pairs, 66% were consistent with the observation for *E. coli* and *S. enterica*, such that variance in selection-driven gene sequence optimisation explained on average 8% of variance in *k*_*s*_ between genes (Fig. 5e). Thus, the extent to which transcript sequences are jointly optimised for cost and efficiency is sufficient to explain a significant component of variation in molecular evolutionary rate between genes within a species. Moreover, selection-driven cost-efficiency optimality is also sufficient to explain the correlation between the rates of synonymous and non-synonymous mutations.

## Discussion

Differences in molecular evolution rates between species are thought to be mainly due to differences in organism generation-time [27]. However, differences in evolutionary rates between genes in the same species lack a complete mechanistic explanation. Prior to the study presented here, it was known that functional constraints of the encoded protein sequence contribute to the constraint of the rate of non-synonymous changes [28]. It had also been observed that mRNA abundance and patterns of codon bias correlated with the evolutionary rate of genes [29,30], and that rates of synonymous and non-synonymous changes were correlated [26]. The study presented here unifies these prior observations and provides a mechanistic explanation for both variation and correlation in molecular evolution rates of genes. Specifically, this study shows that stochastic mutations in gene sequences are more likely to result in deleterious alleles in proportion to the extent to which that gene sequence has been jointly optimised by natural selection for reduced transcript biosynthetic cost and enhanced translational efficiency.

The mechanism provided here also explains the relationship between mRNA abundance and gene evolutionary rate. Specifically, functional constraints on protein abundance stipulate the quantity of mRNA required to produce that protein. The more mRNA that is required, the greater the percentage of total cellular resources that must be invested within that transcript. The mechanism simply entails that the more transcript that is present, the stronger the selective pressure will be to reduce the cellular resources committed to that transcript. Importantly, minimising these resources can be achieved both by using codons that require fewer resources for their biosynthesis, or by utilising translationally efficient codons that increase the protein to transcript ratio and therefore reduce the amount of transcript required to produce the same amount of protein. Overall, this study reveals how the economics of gene production is a critical factor in determining both the evolution and composition of genes.

## Conclusions

Codon use is biased across the tree-of-life, with patterns of bias varying both between species and between genes within the same species. Here we demonstrate that variation in tRNA content between species creates a corresponding variation in the codon cost-efficiency trade-off whereby codons that cost the least to biosynthesise are not equally translationally efficient in all species. We show that irrespective of the codon cost-efficiency trade-off, natural selection performs multi-objective gene sequence optimisation so that transcript sequences are optimised to be both low cost and highly translationally efficient, and that the nature of this trade-off constrains the extent of the solution. We demonstrate that this multi-objective optimisation is dependent on mRNA abundance, such that the transcripts that comprise the largest proportion of cellular mRNA are those that experience the strongest selection to be both low cost and highly efficient. Finally, we show that the extent to which a gene sequence is jointly optimised for reduced transcript cost and enhanced translational efficiency is sufficient to explain a significant proportion of the variation in the rate of gene sequence evolution. Furthermore, it is sufficient to explain the phenomenon that the rate of synonymous and non-synonymous mutation for a gene is correlated [26].

## Methods

### Data sources

1,320 bacterial genomes were obtained from the NCBI (www.ncbi.nlm.nih.gov). In order to avoid over-sampling of more frequently sequenced genera, the number of species from each genus was restricted to 5 with a maximum of 1 species for each genus. Therefore, the 1,320 species sampled in this study were distributed among 730 different genera. Only genes that were longer than 30 nucleotides, had no in-frame stop codons, and began and ended with start and stop codons respectively were analysed. Each species in this analysis contained a minimum of 500 genes that fit these criteria. Full details of species names, genome accession numbers, strain details and selection coefficients are provided in Supplementary Table 1.

### Evaluation of translational efficiency (tAI)

To obtain the number of tRNA genes in each genome, tRNAscan was run on each of the 1,320 bacterial genomes [31]. This current version (1.4) of tRNAscan is unable to distinguish between tRNA-Met and tRNA-Ile with the anticodon CAT. Thus tRNA-Ile(CAT), while present, is not detected in any of the genomes. To compensate for this a single copy of tRNA-Ile with the anticodon CAT was added to the tRNA counts for each species if more than one tRNA-Met(CAT) was found. The tRNA adaptation index (tAI)[21], which considers both the tRNA gene copy number and wobble-base pairing when calculating the translational efficiency of a codon was evaluated using the optimised *s*_*ij*_ values for bacteria obtained by Tuller et al [32] and the equation developed by dos Reis et al [20]. *s*_*uu*_ was set to 0.7 as proposed by Navon et al [33] and *s*_*uc*_ was set to 0.95 as U_34_ has been shown to have weak codon-anticodon coupling with cytosine [34]. Each species in this analysis was able to translate all codons, was not missing key tRNAs and did not require unusual tRNA-modifications.

### Calculation of relative codon cost and efficiency

Codon biosynthetic cost and translational efficiency were calculated relative to other synonymous codons such that the synonymous codon with the greatest value had a relative cost or efficiency of 1. For example, the nitrogen cost of GCC is 11 atoms. The most expensive synonymous codon is GCG/GCA (13 atoms). Therefore the relative cost of GCC is 11/13 = 0.85. The same evaluation was done to calculate codon translational efficiency.

### CodonMuSe: A fast and efficient algorithm for evaluating drivers of codon usage bias

The SK model [2] was used to infer the joint contribution of mutation bias, selection acting on codon biosynthetic cost and selection acting on codon translational efficiency to biased synonymous codon use. To facilitate the large scale comparative application of this model a rapid, stand-alone version was implemented in python.

The algorithm, instructions for use, and example files are available for download at https://github.com/easeward/CodonMuSe. For each species, the values of *M*_*b*_, *S*_*c*_ and *S*_*t*_ were inferred using the complete set of protein coding genes and the tRNA copy number inferred using tRNAscan. Further details about the algorithm can be found in Supplemental File 1.

### Comparing selection acting on codon bias and transcript abundance levels

Transcriptome data for *E. coli* str. K-12 MG1655 were downloaded from NCBI (series GSE15534). The raw data was subject to quantile normalisation and background correction as implemented in the NimbleScan software package, version 2.4.27 [35,36]. The three biological replicates for the logarithmic growth phase were available, however the third replicate was inconsistent with the first two and so was excluded from this analysis. As each gene had multiple probes, the average probe value for each gene was taken. The three-parameter CodonMuSe model using the value for *Mb* estimated from a genome-wide analysis was run for each of the 4099 genes in *E. coli* individually, and thus values for *Sc* and *St* were obtained for each gene. The values for these selection coefficients were plotted against relative mRNA abundance data described above [35].

### Calculating the extent to which gene sequences were jointly optimised for cost and efficiency

To define the extent to which a sequence has been jointly optimised for both biosynthetic cost and translational efficiency the relative Pareto optimality of each gene was calculated. To do this, the boundaries of sequence space were defined as in Supplementary Fig. S2. Here, the cost-efficiency Pareto frontier is the full set of coding sequences that are Pareto efficient, where it is impossible to change the codons of the sequence to make the transcript cheaper without making it less efficient (or vice versa) (red frontier, Supplementary Fig. S2). The opposite frontier is the full set sequences where it is impossible to change the codons of the sequence to make the transcript more expensive without making it more efficient (or vice versa) (blue frontier, Supplementary Fig. S2). Thus, the extent to which transcript sequences were jointly optimised for both biosynthetic cost and translational efficiency was evaluated as the relative distance of a given gene to the cost-efficiency Pareto frontier for the sequence constrained by the amino acid sequence, i.e. 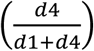 * 100 (Supplementary Fig. S2). Therefore, a value of 100% optimisation represents a gene that lies on the Pareto frontier. Genes that are less than 100% optimised occupy the space between the cost-efficiency Pareto frontier (red frontier) and the opposite frontier (blue frontier, minimising transcript efficiency or maximising cost) for that amino acid sequence (Supplementary Fig. S2).

### Calculation of molecular evolution rates

Molecular evolutionary rates (*k*_*a*_ and *k*_*s*_ values) were calculated for orthologous genes in *E. coli* and *S. enterica*. 2,468 single-copy orthologous genes were identified for *E. coli* and *S. enterica* using OrthoFinder v1.1.4 [37]. These sequences were aligned at the amino acid level using MergeAlign [38] and this alignment was then rethreaded with the coding sequences to create codon-level nucleotide alignments. Only aligned sequences longer than 30 nucleotides with less than 10% gaps were used. Gapped regions were removed and KaKs_Calculator 2.0 [39] was run using the GMYN model to evaluate *k*_*a*_ and *k*_*s*_ values for each pair of aligned nucleotide sequences. As the molecular evolution rates represent the average of the mutation rates of the gene-pair since they last shared a common ancestor, these rates were compared to the average optimality of the same gene-pair in both species.

The same analysis was conducted on 1,066 additional pairs of species obtained by exhaustive pairwise comparison of all species that were within the same genus. These 1,066 pairwise comparisons were filtered to remove those with *k*_*s*_ saturation (i.e. mean *k*_*s*_ > 1) and fewer than 1,000 genes. This filtered set contained 177 species pairs.

### Linear regression analyses

All linear regression analyses were conducted using the lm package in R. In all cases, p-values quoted are the p-values for the linear regression model.

## Declarations

### Ethics approval and consent to participate

Not applicable

### Consent for publication

Not applicable

### Availability of data and material

The datasets generated and/or analysed during the current study are available from the corresponding author on reasonable request.

### Authors’ contributions

SK and EAS conceived the study, EAS conducted the analysis, EAS and SK wrote the manuscript. Both authors read and approved the final manuscript.

## Acknowledgements

Not applicable

---

**Supplementary Figure 1. Correlation between tAI and codon biosynthetic cost with *K*_*s*_ and *K*_*a*_ for *Escherichia coli* and *Salmonella enterica*.**

**a)** Scatter-plot of log_10_(*K*_*S*_) compared to average tAI per codon per gene (y = −0.3x + 2.2, R^2^ = 0.25). **b)** Scatter-plot of log10(*K*_*a*_) compared to average tAI per codon per gene (y = −0.3x + 1.8, R^2^ = 0.26). **c)** Scatter-plot of log_10_(*K*_*s*_) compared to average cost per codon per gene (y = −0.1x + 11.4, R^2^ = 0.02). **d)** Scatter-plot of log_10_(*K*_*a*_) compared to average cost per codon per gene (y = −0.1x + 11.3, R^2^ = 0.01).

**Supplementary Figure 2. Example cost-efficiency Pareto frontier for a short amino acid sequence**.

**a)** Scatter plot of the 64 possible coding sequences encoding the amino acid sequence MTGCD. Red dots indicate coding sequences that are positioned on the best cost-efficiency Pareto frontier (the least expensive, most translationally efficient sequences possible). Blue dots indicate coding sequences that are positioned on the worst cost-efficiency Pareto frontier (the most expensive, least translationally efficient sequences possible). **b)** Evaluating the cost-efficiency optimality of a coding sequence. d1 is the minimum distance between a given coding sequence and the best cost-efficiency Pareto frontier (red) for that amino acid sequence. d4 is the minimum distance of the same gene to the worse cost-efficiency Pareto frontier for that amino acid sequence (blue). The percent optimality of the coding sequence is evaluated as 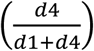 * 100.

## References

1. Farmer IS, Jones CW. The Energetics of Escherichia coli during Aerobic Growth in Continuous Culture. Eur. J. Biochem. 1976;67:115–22.

2. Seward EA, Kelly S. Dietary nitrogen alters codon bias and genome composition in parasitic microorganisms. Genome Biol. 2016;17:1–15.

3. Chen W-H, Lu G, Bork P, Hu S, Lercher M. Energy efficiency trade-offs drive nucleotide usage in transcribed regions. Nat. comminications. 2016;7:1–10.

4. Horn D. Codon usage suggests that translational selection has a major impact on protein expression in trypanosomatids. BMC Genomics. 2008;9:1–11.

5. Rocha EPC. Codon usage bias from tRNA’s point of view: Redundancy, specialization, and efficient decoding for translation optimization. Genome Res. 2004;14:2279–86.

6. Sorensen M a, Kurland CG, Pedersen S. Codon usage determines translation rate in Escherichia coli. J. Mol. Biol. 1989;207:365–77.

7. Hu H, Gao J, He J, Yu B, Zheng P, Huang Z, et al. Codon Optimization Significantly Improves the Expression Level of a Keratinase Gene in Pichia pastoris. PLoS One. 2013;8:e58393.

8. Akashi H. Synonymous codon usage in Drosophila melanogaster: Natural selection and translational accuracy. Genetics. 1994;136:927–35.

9. Shah P, Gilchrist M a. Explaining complex codon usage patterns with selection for translational efficiency, mutation bias, and genetic drift. Proc. Natl. Acad. Sci. U. S. A. 2011;108:10231–6.

10. Precup J, Parker J. Missense misreading of asparagine codons as a function of codon identity and context. J. Biol. Chem. 1987;262:11351–5.

11. Lao PJ, Forsdyke DR. Thermophilic bacteria strictly obey Szybalski’s transcription direction rule and politely purine-load RNAs with both adenine and guanine. Genome Res. 2000;10:228–36.

12. Paz A, Mester D, Baca I, Nevo E, Korol A. Adaptive role of increased frequency of polypurine tracts in mRNA sequences of thermophilic prokaryotes. Proc. Natl. Acad. Sci. U. S. A. 2004;101:2951–6.

13. Goodman DB, Church GM, Kosuri S. Causes and Effects of N-Terminal Codon Bias in Bacterial Genes. Science. 2013;342:475–80.

14. Eskesen ST, Eskesen FN, Ruvinsky A. Natural selection affects frequencies of AG and GT dinucleotides at the 5’ and 3’ ends of exons. Genetics. 2004;167:543–50.

15. Novoa EM, Ribas de Pouplana L. Speeding with control: Codon usage, tRNAs, and ribosomes. Trends Genet. 2012;28:574–81.

16. Zhang F, Saha S, Shabalina SA, Kashina A. Differential Arginylation of Actin Isoforms Is Regulated by Coding Sequence-Dependent Degradation. Science. 2010;329:1534–7.

17. Drummond DA, Wilke CO. Mistranslation-induced protein misfolding as a dominant constraint on coding-sequence evolution. Cell. 2008;134:341–52.

18. Grosjean H, de Crécy-Lagard V, Marck C. Deciphering synonymous codons in the three domains of life: Co-evolution with specific tRNA modification enzymes. FEBS Lett. Federation of European Biochemical Societies; 2010;584:252–64.

19. Brocchieri L. The GC content of bacterial genomes. Phylogenetics Evol. Biol. 2013;1:e106.

20. dos Reis M, Savva R, Wernisch L. Solving the riddle of codon usage preferences: A test for translational selection. Nucleic Acids Res. 2004;32:5036–44.

21. dos Reis M, Wernisch L, Savva R. Unexpected correlations between gene expression and codon usage bias from microarray data for the whole Escherichia coli K-12 genome. Nucleic Acids Res. 2003;31:6976–85.

22. Drummond DA, Wilke CO. The evolutionary consequences of erroneous protein synthesis. Nat Rev Genet. 2009;10:715–24.

23. Ran W, Higgs PG. Contributions of Speed and Accuracy to Translational Selection in Bacteria. PLoS One. 2012;7:1–7.

24. Pal C, Papp B, Hurst LD. Highly expressed genes in yeast evolve slowly. Genetics. 2001;158:931.

25. Brandis G, Hughes D. The Selective Advantage of Synonymous Codon Usage Bias in Salmonella. PLOS Genet. 2016;12:e1005926.

26. Sharp PM. Determinants of DNA sequence divergence between Escherichia coli and Salmonella typhimurium: Codon usage, map position, and concerted evolution. J. Mol. Evol. 1991;33:23–33.

27. Weller C, Wu M. A generation-time effect on the rate of molecular evolution in bacteria. Evolution (N. Y). 2015;69:643–52.

28. Zuckerkandl E. Evolutionary processes and evolutionary noise at the molecular level. I. Functional Density in Proteins. J. Mol. Evol. 1976;7:167–83.

29. Sharp PM, Li WH. The rate of synonymous substitution in enterobacterial genes is inversely related to codon usage bias. Mol. Biol. Evol. 1987;4:222–30.

30. Drummond DA, Raval A, Wilke CO. A single determinant dominates the rate of yeast protein evolution. Mol. Biol. Evol. 2006;23:327–37.

31. Schattner P, Brooks AN, Lowe TM. The tRNAscan-SE, snoscan and snoGPS web servers for the detection of tRNAs and snoRNAs. Nucleic Acids Res. 2005;33:686–9.

32. Sabi R, Tuller T. Modelling the Efficiency of Codon–tRNA Interactions Based on Codon Usage Bias. DN Res. 2014;21:511–25.

33. Navon S, Pilpel Y. The role of codon selection in regulation of translation efficiency deduced from synthetic libraries. Genome Biol. 2011;12:1–10.

34. Näsvall SJ, Chen P, Björk GR. The modified wobble nucleoside uridine-5-oxyacetic acid in tRNA Pro cmo 5 UGG promotes reading of all four proline codons in vivo The modified wobble nucleoside uridine-5-oxyacetic acid in tRNA Pro cmo UGG promotes reading of all four proline codons in vi. 2004;10:1662–73.

35. Cho B-K, Zengler K, Qiu Y, Park YS, Knight EM, Barrett CL, et al. Elucidation of the transcription unit architecture of the Escherichia coli K-12 MG1655 genome. Nat. Biotechnol. Nature Publishing Group; 2009;27:1043–9.

36. Bolstad BM, Irizarry RA, Astrand M, Speed TP. A comparison of normalization methods for high density oligonucleotide array data based on variance and bias. Bioinformatics. 2003;19:185–93.

37. Emms DM, Kelly S. OrthoFinder: solving fundamental biases in whole genome comparisons dramatically improves orthogroup inference accuracy. Genome Biol. Genome Biology; 2015;16:1–14.

38. Collingridge PW, Kelly S. MergeAlign: improving multiple sequence alignment performance by dynamic reconstruction of consensus multiple sequence alignments. BMC Bioinformatics. 2012;13:1–10.

39. Wang D, Zhang Y, Zhang Z, Zhu J, Yu J. KaKs_Calculator 2.0: A Toolkit Incorporating Gamma-Series Methods and Sliding Window Strategies. Genomics, Proteomics Bioinforma. Beijing Institute of Genomics; 2010;8:77–80.

